# A CMOS-based highly scalable flexible neural electrode interface

**DOI:** 10.1101/2022.11.03.514455

**Authors:** Eric T. Zhao, Jacob Hull, Nofar Mintz Hemed, Hasan Uluşan, Julian Bartram, Anqi Zhang, Pingyu Wang, Albert Pham, Silvia Ronchi, John R. Huguenard, Andreas Hierlemann, Nicholas A. Melosh

## Abstract

Perception, thoughts, and actions are encoded by the coordinated activity of large neuronal populations spread over large areas. Using thin film electrocorticography (ECoG) arrays, this cortical activity has been used to decode speech and individual finger movements, enabling neuroprosthetics, and to localize epileptic foci. However, the connectorization of these multi-thousand channel thin-film arrays to external circuitry is challenging; current state-of-the-art methods are complex, bulky, and unscalable. We address this shortcoming by developing an electrode connector based on an ultra-conformable thin film electrode array that self-assembles onto hard silicon chip sensors, such as microelectrode arrays (MEAs) or camera sensors enabling large channel counts at high density. The interconnects are formed using microfabricated electrode pads suspended by thin support arms, termed flex2chip. Capillary-assisted assembly drives the pads to deform towards the chip surface, and van der Waals forces maintain this deformation, establishing mechanical and Ohmic contact onto individual pixels. We demonstrate a 2200-channel array with a channel density of 272 channels / mm^2^ connected to the MEA through the flex2chip interconnection method. Thin film electrode arrays connected through the flex2chip successfully measured extracellular action potentials ex vivo. Furthermore, in a transgenic mouse model for absence epilepsy, *Scn8a*^+/-^, we observed highly variable propagation trajectories at micrometer scales, even across the duration of a single spike-and-wave discharge (SWD).

## Introduction

Perception, thoughts, and actions involve the coordinated activity of large populations of neurons in multiple regions of the brain (*1*–*3*). Non-penetrative, subdural ECoG grids laid on top of the brain surface are the gold standard for recording population-level activity, measured from the local field potential (LFP). Neurophysiological recordings with ECoG grids have been successfully used for speech synthesis (*4, 5*), reproduction of arm movements (*6*), and spatial localization of ictal onset zones (*7*). They have also been used to characterize cortical travelling waves (*8*) which have been shown to modulate perceptual sensitivity (*9*). As such, ECoG grids are a favorable modality for brain-computer interface (BCI) applications, localization of epileptic foci for clinical epilepsy diagnosis and targeted tissue resection, and as a basic science tool.

Ultra-conformable thin film flexible devices that can conform to the curvilinear surface of the brain are a promising technology to capture cortex-wide spatiotemporal dynamics (*10*– *12*). Microfabrication and advanced lithography methods have enabled the creation of thin-film arrays with hundreds to thousands of recording sites (*13, 14*). However, the key bottleneck lies not in the fabrication of such devices but in the connectorization between each electrode and the external circuitry. Due to the bulkiness of existing connectorization methods such as wire-bonding, anisotropic conductive film (*15*), and ultrasonic-on-bump bonding (*16*), current implementations of multi-thousand channel count, passive thin-film neural interfaces are highly complex, bulky, and unscalable. Examples include the stacking of 16 application-specific integrated circuits (ASICs) on 8 circuit boards to achieve 1024 channels (*17, 18*), modularization of 12 ASICs on a single circuit board for 3072 channels (*19*), and the use of 2 CPU sockets, 2 circuit boards and 32 ASICs for 2048 channels (*11*).

Consequently, the field has focused on adapting this technology for active, multiplexed thin-film arrays that monolithically integrate the recording electrodes with amplifiers and analog-to-digital converters, bypassing the one electrode per input/output (I/O) limit (*12, 20*). However, these active thin-film arrays suffer from increased noise, sub-kilohertz sampling rates, lower channel counts, and significantly increased device size due to the difficulty in fabricating high mobility, high mechanical flexibility, and small-size flexible transistors (*21*). On the other hand, silicon-based large-scale complementary metal-oxide semiconductor microelectrode arrays (CMOS-MEAs) have built on decades of development in active pixel sensors, traditionally used in cameras, to provide excellent signal-to-noise ratio, high sampling rates, and scalability up to tens of thousands of channels at high densities (*22*–*24*). However, their rigid form factor is incompatible for interfacing with the brain surface.

Here we sought to combine the scalability and exceptional performance of CMOS-MEAs with the ultra-conformable form factor of flexible devices. We achieve this by developing a scalable, high-density connectorization strategy which can form thousands of interconnections between the electrode pads on the flexible device and the pixels on the CMOS-MEA at a high density. The flexible device extends out from the CMOS-MEA through long leads, converging at the distal end as an array that interfaces with the brain (Fig. 1A).

**Figure 1.**
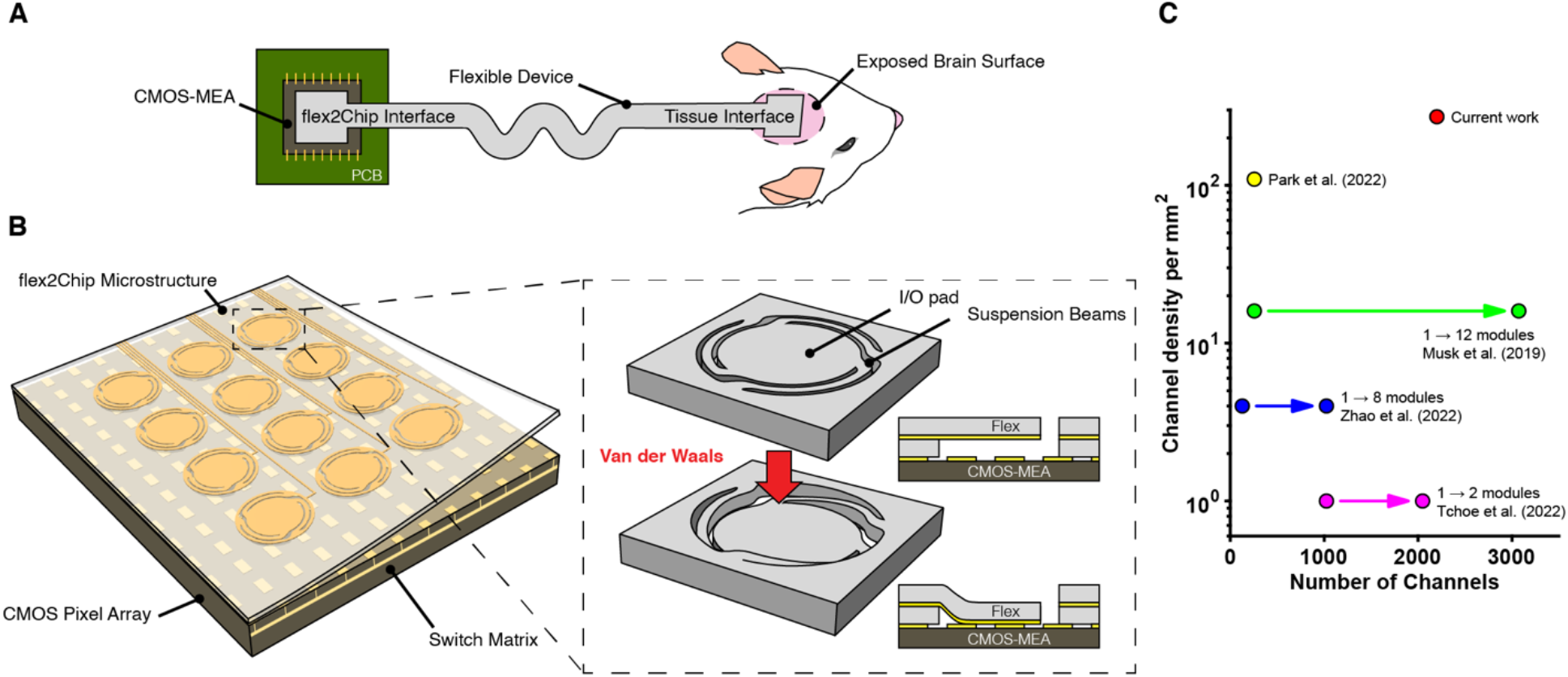
flex2chip design. (**A**) A schematic of the flexible device. At the chip interface, flex2chip microstructures enable multi-thousand Ohmic interconnections to a CMOS-MEA. At the tissue interface, the device terminates as an electrode array for high density recording. (**B**) The flex2chip interface consists of an array of deformable flex2chip microstructures, integrated with the underlying CMOS-MEA. The flex2chip microstructures consist of three suspended beams and an I/O pad. Capillary forces deform the I/O pad to contact the CMOS pixels, upon which Van der Waals forces become significant to establish structural and electrical contact. (**C**) Current approaches to multi-thousand channel counts rely on the modularization of multiple ASICs and circuit boards. The flex2chip structures facilitate 2200 individual Ohmic connections to a single ASIC in a 3.5 mm x 2.1 mm area, yielding a channel density of 272 channels per mm^2^.

We design a microstructure, termed flex2chip, which consists of microfabricated 1-μm-thick and 2-μm-wide supporting arms suspending each individual electrode pad. We leverage the ultra-compliant nature of the supporting arms to allow capillary and van der Waals forces to deform the pad to establish mechanical and electrical contact with the underlying pixels (Fig. 1B). This flex2chip interconnection is self-assembled and does not require any equipment or lithographic post-processing to facilitate electrical connectivity. We demonstrate a 2200-channel device with a connection interface area of 3.85 × 2.1 mm^2^, 17 times denser than that of current multi-thousand channel devices (Fig. 1C). Importantly, our method could achieve multi-thousand channel counts without relying on the modularization of multiple ASICs and circuit boards.

We further show the utility of our devices in neuroscience and biologically relevant applications. We validate that our device can record extracellular action potentials with a high signal-to-noise ratio in acute ex vivo cerebellar slices. Finally, we demonstrate the efficacy of our device and its potential clinical utility through the detection and characterization of sub-millimeter traveling waves during absence seizures on the cortical surface of awake and behaving epileptic mice. Here, our high-density flexible probes combined with the temporal resolution of the CMOS-MEA enabled the observation of highly variable propagation trajectories at micrometer scales even across the duration of a single SWD.

## Results

### System design and connectorization for the flex2chip devices and microstructures

The flex2chip devices were fabricated using standard microfabrication procedures (Materials and Methods). The devices consist of a 2200 channel array of 1-μm-wide and 100-nm-thick platinum leads at a 2 μm pitch, sandwiched between two 1-μm-thick polyimide sheets that serve as substrate and dielectric layers (Fig. 1A). At the flex2chip interface, these leads branch out to an array of I/O pads which interface with the CMOS-MEA (Fig. 1B, Fig. S1A). At the tissue interface, the leads branch out into an array of recording/stimulation pads (Figs. S1B and C).

At the flex2chip interface, the dielectric layer is etched so that the conductive I/O pad is exposed. However, the pad is recessed by 1 μm and unable to directly contact external circuitry. Unlike previous approaches which rely on the addition of external components, e.g., gold wires for wire bonding and microparticles for anisotropic conductive films, we instead modified the I/O pad itself to facilitate connectorization.

Here, we developed a microstructure on the I/O pad, termed flex2chip, enabling the pad to deform downwards to mate with contact pads on external circuitry. The flex2chip consists of a 35-μm-diameter I/O pad suspended by three suspension beams, 2 μm wide and 10 μm long (Fig. 1B inset top, Fig. S2). The conductive leads run from the I/O pad, across the suspension beams, and onto the main substrate.

The low bending stiffness and flexibility of the suspension beams permit the I/O pad to deform relative to the flexible device and toward the underlying pixel on the CMOS array under capillary forces (see the section below) until the platinum electrodes make physical and electrical contact with the gold-plated pixels, upon which van der Waals forces hold the microstructure in its collapsed configuration (Fig. 1B, inset bottom).

We included three suspension beams which mechanically prevent the pad from twisting or flipping over during handling and assembly. The interelectrode pitch was set at 50 μm (Figs. 2A and B, Fig. S2D), greater than the pixel pitch, so that adjacent pads will not be shorted (*22, 25, 26*). This simplifies assembly, allowing rotational and translational degrees of freedom.

**Figure 2.**
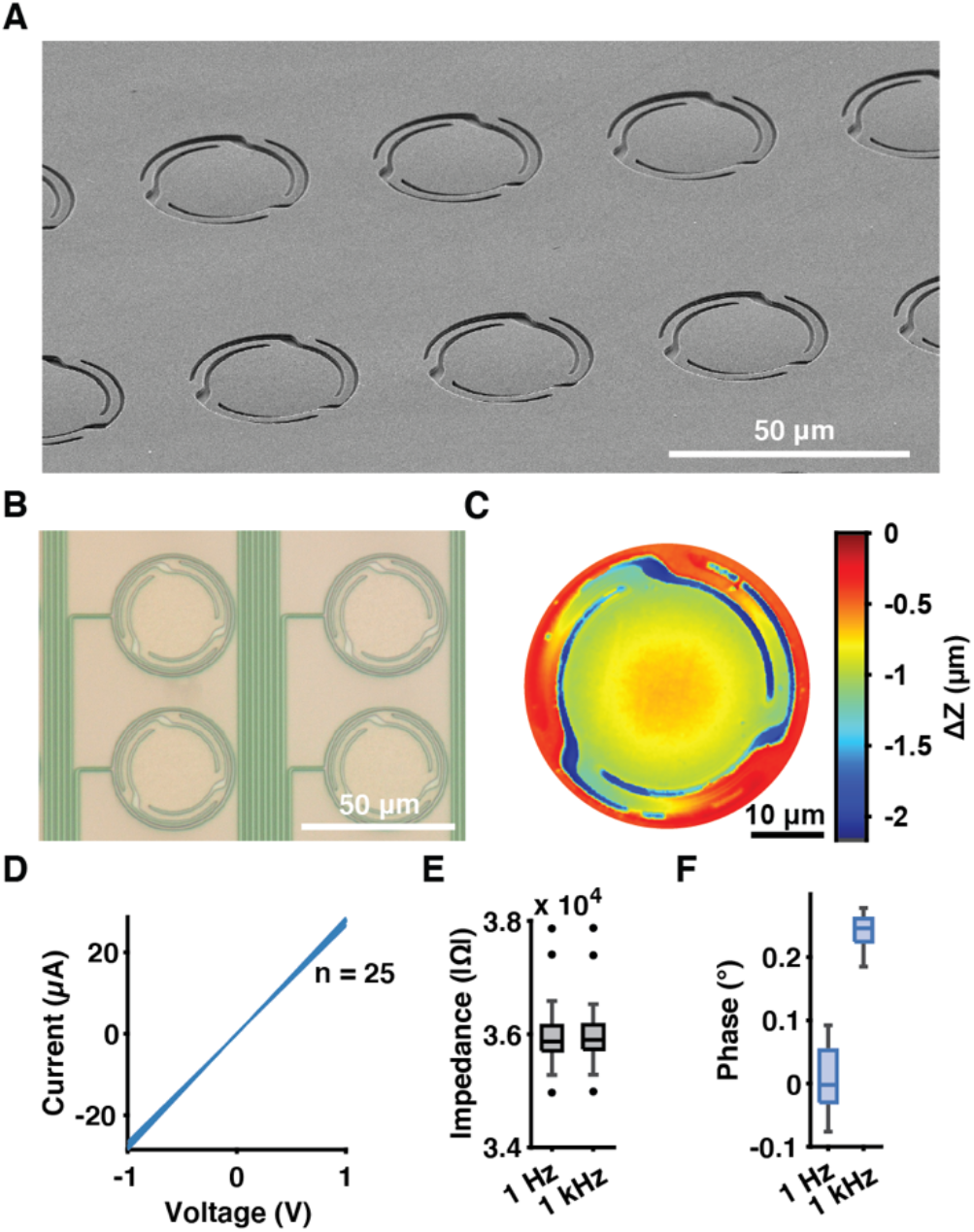
flex2chip structure and electrical characteristics. (**A**) A SEM image of the flex2chip device laid on top of a planar glass surface. A slight deformation of the 40-µm-diameter I/O pad towards the underlying surface enabled by the suspension beams can be observed. (**B**) Each flex2chip microstructure is routed by a 1-µm-wide trace towards the distal end of the device, terminated by a recording/stimulation pad. (**C**) The optical profilometer measurement confirms that the flex2chip I/O pad has fully deformed by 1 µm to contact the underlying surface, and that van der Waals forces are sufficient to hold the structure in its deformed configuration, which facilitates stable mechanical contact. (**D**) The mechanical contact is Ohmic, as seen in the resistive behavior of the IV curve (n = 25). (**E**) The impedance magnitude of the flex2chip interface is 36.0 ± 0.60 and 36.0 ± 0.59 kΩ at 1 and 1000 Hz respectively (n = 25). (**F**) The phase of the flex2chip interface is 0.01 ± 0.05° and 0.24 ± 0.02° at 1 and 1000 Hz respectively (n = 25).

At the tissue interface, the leads fanout to an array of recording/stimulation pads (Fig. S1C). The distal end geometry can be freely customized for the specific biological system of interest. For example, we can have a contiguous sheet for ECoG grids or shanks for intracortical insertion (*18*).

### Microstructure deformation assisted by capillary assembly

We leverage the flexibility of the flex2chip microstructures to establish connectorization between the I/O pad and CMOS-MEA pixel. This deformation is achieved under the action of capillary and van der Waals forces, which operate at micrometer and nanometer length scales, respectively. When the flexible device is placed on the CMOS-MEA the I/O pad is 1 μm away from the underlying pixel (the thickness of the dielectric layer) (Fig. 1B inset top). We initiate the microstructure deformation by applying a thin layer of isopropyl alcohol (IPA) between the device and CMOS-MEA. The capillary force of the liquid bridge formed between the I/O pad and the pixel ‘pulls’ the pad towards the pixel as the solvent evaporates. After the IPA fully evaporates and the pad reaches its fully collapsed structure van der Waals forces become significant enough to hold the microstructure in its collapsed configuration (Fig. 1B inset bottom).

We characterized this deflection by assembling a device on a CMOS-MEA phantom. Optical profilometry of the flex2chip microstructures showed that the structures have fully deformed and contacted the underlying chip (Fig. 2C). The deformation was observed to be uniformly distributed across the three suspension beams and the I/O pad was uniformly displaced by 1 μm relative to the bulk device. The center of the I/O pad was slightly curved upward, indicative of residual stress from the device fabrication. Scanning electron microscopy (SEM) images also support the optical profilometer results, with a slight deformation visible despite its 1:40 aspect ratio (Fig. 2A).

The electrical characteristics of the flex2chip microstructure interface were evaluated to assess the quality of electrical contact and whether stray capacitance was introduced which may affect the quality of the recording, electrical stimulation, or electrochemical measurements. Here, we shorted the distal end of the flex2chip device to a Gallium droplet as a liquid contact. The current-voltage characteristic curve (I-V) exhibits a clear linear relationship, characteristic of Ohmic resistors, with a resistance of 36.0 ± 0.60 kΩ (Fig. 2D, Fig. S3A). Furthermore, impedance spectroscopy indicated an Ohmic connection between the microstructure and chip, with an average phase of 0.01 ± 0.05° and 0.24 ± 0.02° at 1 and 1000 Hz, respectively (Fig. 2E and F, Fig. S3B). This finding confirmed that there is no capacitive impedance at the physiological frequencies of interest. Impedance contributions included cumulative contributions from the flex2chip interface and trace resistances but were insignificant compared to electrode-electrolyte impedance.

### Electrical performance through CMOS-MEA

Having confirmed the quality of the mechanical and electrical contact of our flex2chip structures, we sought to characterize the electrode yield and recording quality of our device using a CMOS-MEA recording chip. Here, we fabricated a 720- and 2200-channel flex2chip array (Fig. S1A and Fig. S4A). The I/O pads are distributed across a 3.85 mm x 2.10 mm area, the active dimensions of the CMOS-MEA (Maxwell Biosystems Inc., Zurich, Switzerland), which has 26,400 active pixel sensors at a pitch of 17.5 μm. As our interelectrode pitch is greater than the pixel pitch, no alignment is needed during assembly.

We evaluated the connectivity yield of the device by applying a sinuisoidal waveform across all the electrodes. we submerged the distal end of the device (Figs. S1B and C, Figs. S4B and C) in a phosphate buffered saline (PBS) bath along with a Pt reference electrode. We applied a sinusoidal waveform at 1 kHz at the reference electrode and scanned the response of each of the 26,400 active pixel sensors. The pixels that were connected to our flex2chip microstructure subsequently then recorded the resulting waveform (Fig. 3A). By cross-referencing with the geometry of the flexible array (Fig. 3B) the position of each microstructure on the CMOS-MEA was localized. As the CMOS-MEA can only measure from 1024 channels simultaneously, we programmed the switch matrix to select only the pixels that corresponded to the individual microstructures to record from.

**Figure 3.**
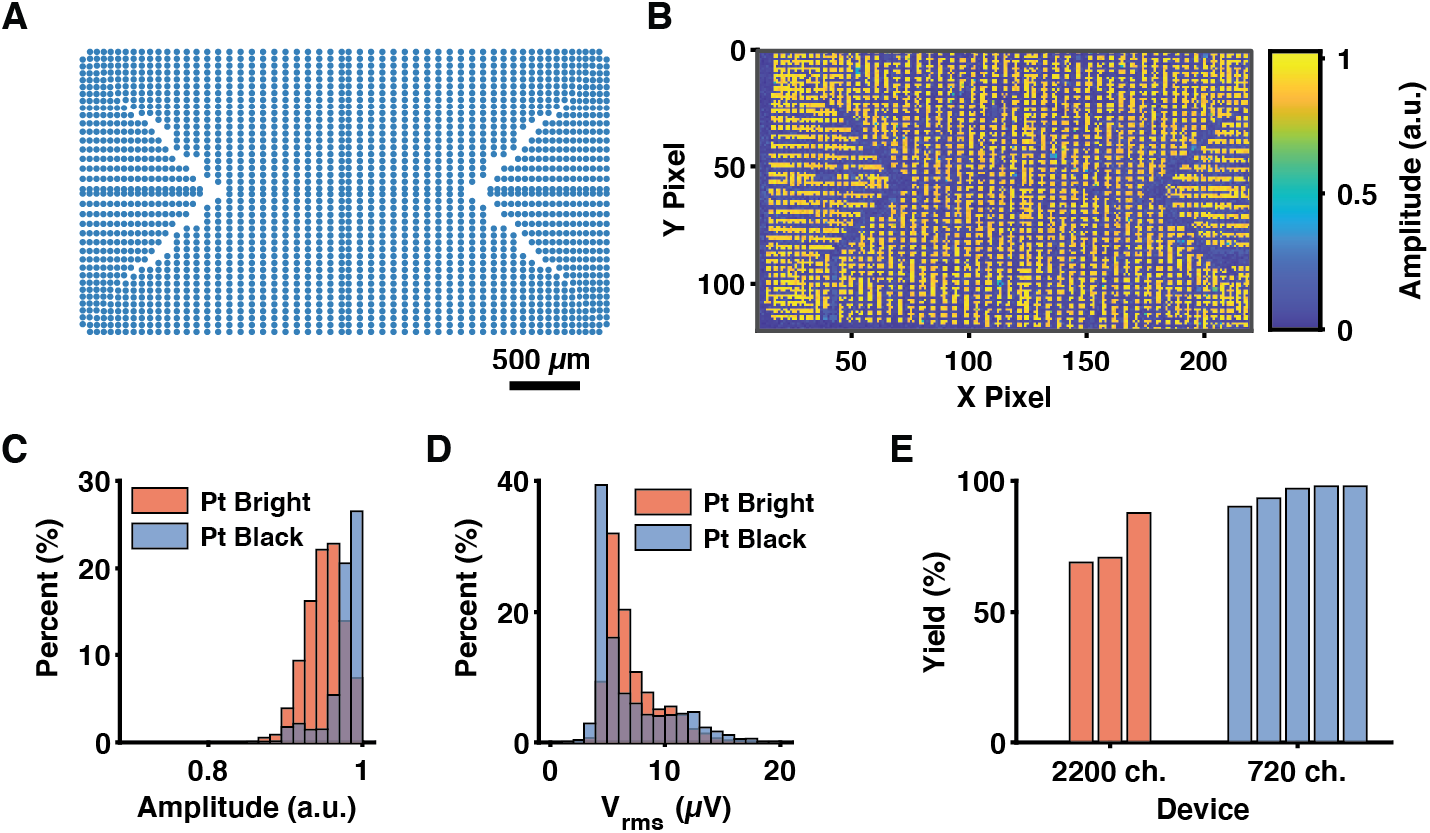
Connectivity to the CMOS-MEA. (**A**) The flex2chip pad layout. The traces are routed out on all four sides to maximize the pad density interfacing with the CMOS-MEA as seen in Fig. S4A. (**B**) The heatmap of the relative signal amplitude measured by each pixel in the CMOS-MEA. The signal is generated through the reference electrode in a saline bath by an external function generator. (**C**) The histogram of the relative signal amplitude of the pixels connected to the flex2chip microstructures. Bright platinum electrodes detected 0.95 ± 0.03 and platinum black-coated electrodes detected 0.99 ± 0.03 with respect to the full-scale injected signal, indicative of minimal attenuation. (**D**) Histogram of the RMS noise was 7.13 ± 2.33 µV for bright platinum and 6.93 ± 3.31 µV for platinum black electrodes. (**E**) The average connectivity yield of the 2200-channel device and 720-channel device was 75.7% and 95.3% respectively.

We then deposited platinum black (Pt black) on the electrode pads to reduce the signal attenuation, Johnson-Nyquist noise, and interelectrode crosstalk (*27*). Here, we leveraged the stimulation units of the CMOS-MEA for voltage controlled electrochemical deposition at -0.5 V at the working electrode for 40 seconds. With bare platinum (Pt bright), the pixels detected 0.95 ± 0.03 of the injected sine wave signal a 5% attenuation. This was further reduced by Pt black, which detected 0.99 ± 0.03 with respect to the full-scale injected signal (Fig. 3C). The root mean squared (RMS) noise was also low, with platinum bright 7.13 ± 2.33 µV and platinum black 6.93 ± 3.31 µV, minimally higher than that of the bare pixels 5.00 ± 1.50 µV (*25*).

The developed connectorization methodology is also highly reliable with an average connectivity yield of 95.3 ± 3.42 % and 75.7 ± 10.4 % for a 720-channel and a 2200-channel device, respectively. The ultra-low footprint of the device is highlighted with a 720-channel device for in vivo recordings as shown in Fig. S2C. We also assessed the stability of the interconnection in an incubator over a month (37 °C, 97% humidity, Heracell 150i, Thermo Fischer Scientific Inc., MA, USA). The yield is virtually unchanged, decreasing from 709 to 705 out of 720 channels (Fig. S3D), with no decrease in the sensitivity (0.99 ± 0.03) (Fig. S3E).

### Multi-site brain slice electrophysiology

Having prepared our flex2chip device, we then tested its ability to record neural activity in an ex vivo preparation of an acute cerebellar brain slice. Here, we used a 720-channel device, where the recording end covers a 3.6 mm x 1.62 mm area with 90 µm pitch (30 × 24 array) with 20-µm-diameter electrode pads (Figs. S4C and D). The dimensions and channel count of the device was chosen to match the size of the tissue slice. The 720-channel device was attached to a 6 mm diameter stainless steel ring and the weighted construct was placed on top of the slice (Fig. S5). The device was fenestrated with 60 μm x 80 μm holes between each electrode to allow sufficient nutrient and oxygen diffusion to keep the slice healthy for the duration of the experiment. As seen in the bright-field microscopy image in Fig. 4A, the neurons were on the same focal plane as the device, indicating good contact.

**Figure 4.**
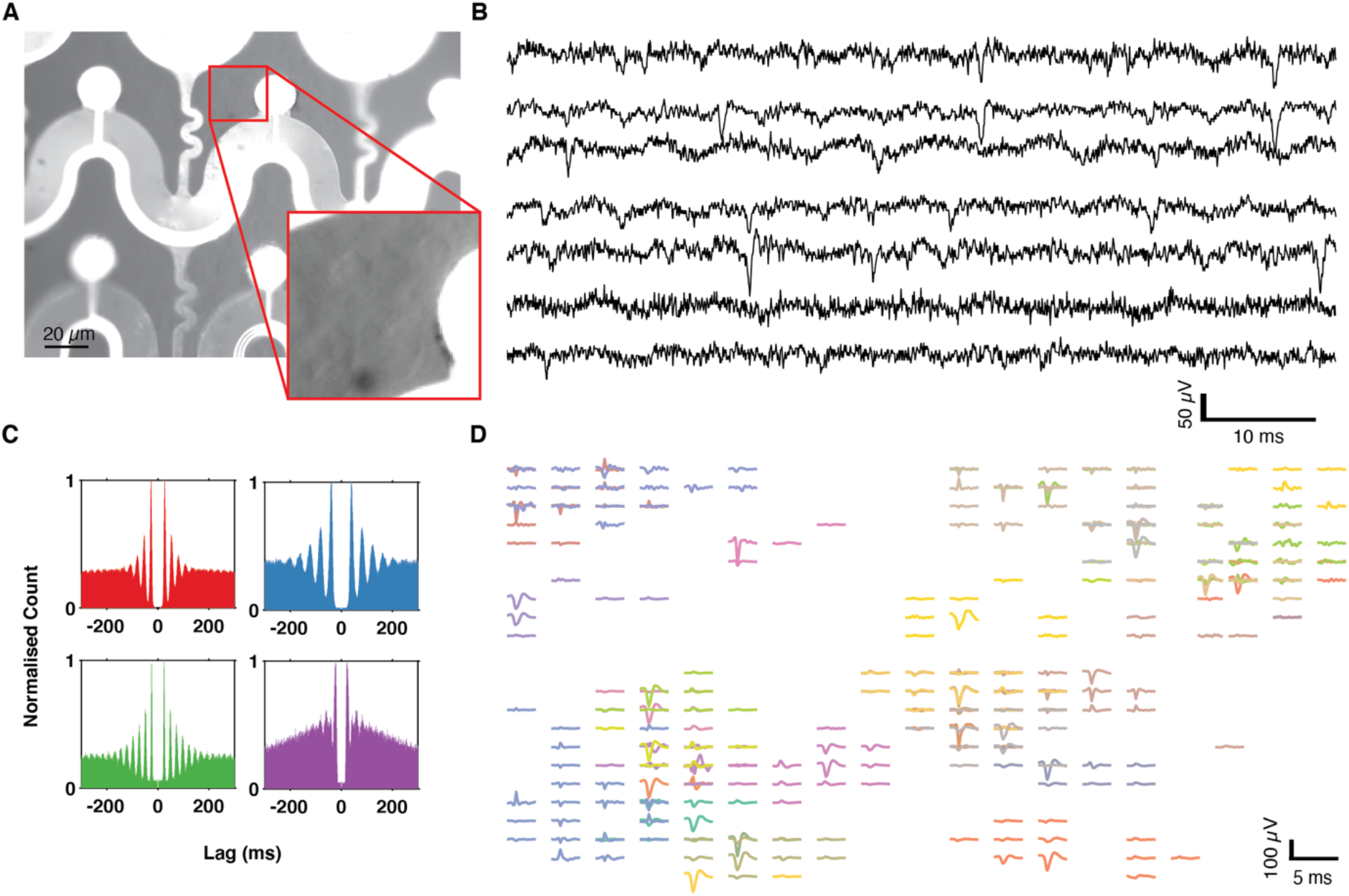
In vitro cerebellar slice recordings. (**A**) Microscopy of the fenestrated 720-channel device. The device and the cerebellar cells were in the same focal plane. The inset has adjusted brightness and contrast highlighting the cerebellar cells. (**B**) Representative traces off the 720-channel device with a 50 Hz notch filter. (**C**) Spike autocorrelograms of cerebellar cells exhibiting regular firing behavior, spanning ± 300 ms and a bin size of 5 ms. The peaks show that the spiking activity occurred at a regular period characteristic of Purkinje cells. (**D**) The spike-triggered average waveform of individually sorted units plotted spatially across the electrodes.

We observed spontaneous spiking activity and isolated 36 individual units from various spatial locations of the slice (Fig. 4B and Fig. 4C). Units were well isolated as indicated by inter-spike interval (ISI) violations within the refractory period (±1.5 ms) below 0.1. The clearly defined peaks in the autocorrelograms demonstrate regular firing behavior typical of cerebellar Purkinje cells (*28*). With an electrode pitch of 90 μm we successfully detected the same neuron across multiple electrodes so that we were able to establish a neural footprint, that is, the spike-triggered average electrical potential distribution across electrodes for a specific unit (Fig. 4D). For multiple neurons, we detected both negative and positive amplitude spikes, indicative of the signal originating from the axon initial segment (negative), and dendritic branches (positive) (*29*). This system thus provides a simple, small form-factor method of recording from hundreds to thousands of electrode sites while retaining high-quality recordings and single-spike level sensitivity.

### Seizure recordings in awake and behaving mice

We next evaluated the performance of the device in a real-world application as an ECoG grid to track the dynamics of seizure propagation with micrometer precision by placing it on the cortical surface of an epileptic mouse (Fig. 5A). Here, we fabricated a 504-channel device with an active area of 760 μm x 760 μm with 20-µm-diameter electrode pads (Fig. 5B and Fig. S4B), customized to record from a 2-mm-diameter craniotomy window above the sensorimotor cortex. The total channel count in this case was constrained simply by the size of the craniotomy. To differentiate local activity at each recording site from volume conducted signals, we utilized the analytic signal method which can identify localized instantaneous frequency and phase information (Fig. 5C). The recordings were performed acutely within 2 hours of the device placement onto the exposed motor and somatosensory cortex. Mice were allowed to run voluntarily on a cylindrical treadmill in a head-strained condition while recording with the device and CMOS-MEA.

**Figure 5.**
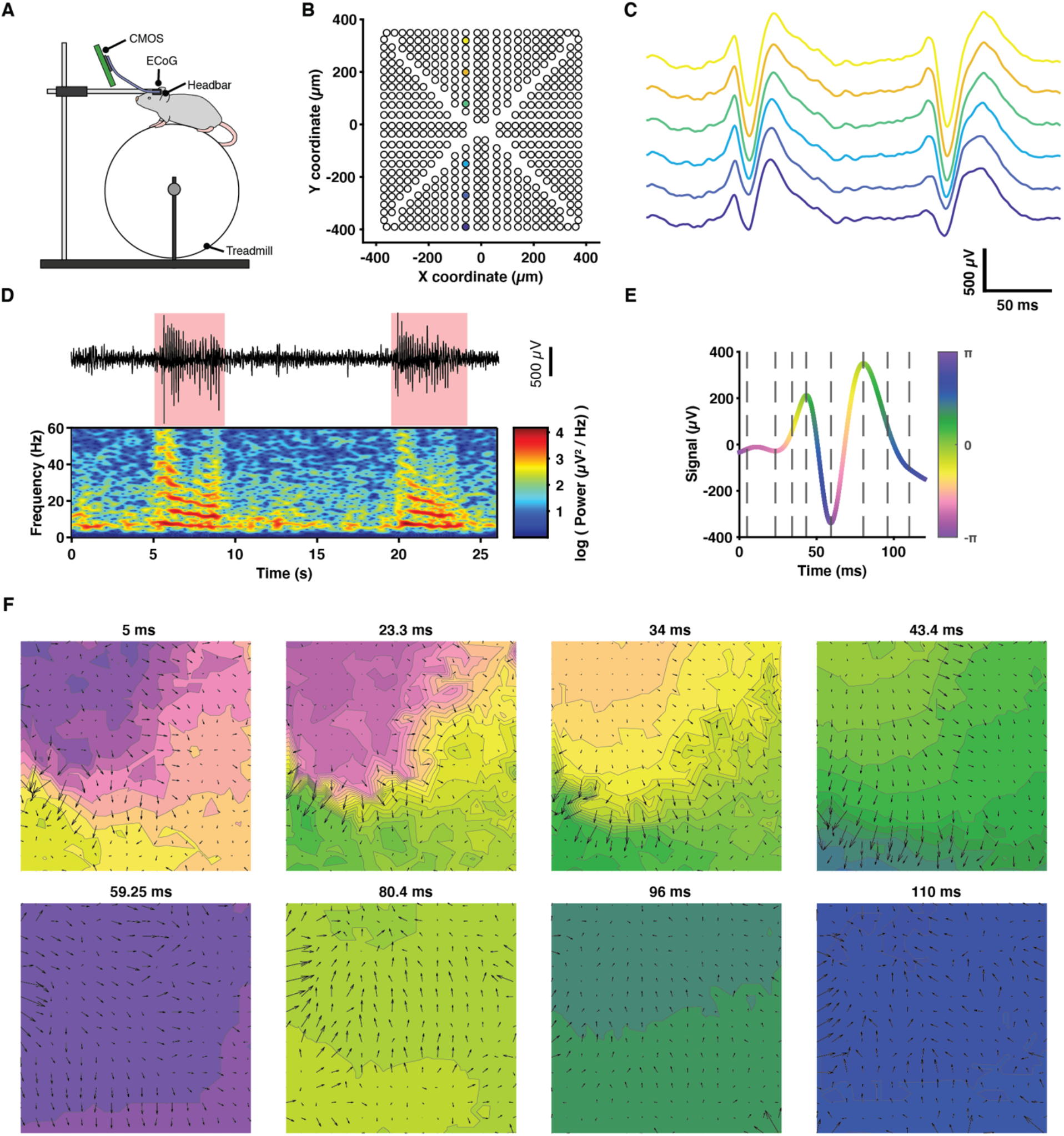
In vivo recording in an awake and moving mouse. (**A**) A schematic of a head-fixed epileptic mouse on a treadmill. A 2-mm-diameter craniotomy exposes the cortex upon which a 504-channel ECoG array spanning 0.76 mm x 0.76 mm is laid. (**B** and **C**) The ECoG electrode layout and representative traces showing SWDs during a seizure, color-coded to match their respective positions. (**D**) (top) A representative trace of a single channel with two seizures highlighted in red. (bottom) The corresponding spectrogram showing the decrease in the frequency band across the duration of the seizure, characteristic of absence epilepsy. (**E**) Representative trace color coded according to the instantaneous phase of the signal. (**F**) Heatmap of the instantaneous phase across the electrode arrays during a single spike wave discharge. Each frame corresponds to a dotted line in Fig. 3E.

The transgenic mouse model *Scn8a*^+/-^ exhibits spontaneous absence epilepsy, characterized by brief periods of unconsciousness and lapse in motor function that have a distinct 7 Hz SWD (*30*). Our single-channel measurements showed the stereotypical 7 Hz SWD with a shift toward low-frequency band power across the duration of a single seizure (Fig. 5D).

We computed the instantaneous phase of the LFP signal to characterize the spatiotemporal propagating wave dynamics that occurs during seizures (*31*). Here, we used the generalized phase (GP), an updated approach of the analytic signal to enable the analysis of wideband signals (*9*). We chose this approach to sufficiently capture the multiple frequency bands associated with the SWD as shown in the spectrogram in Fig. 5D. In short, the GP captures the phase of the largest fluctuation on the recording electrode at any moment in time without distortions due to large low-frequency intrusions or smaller high-frequency inclusions (Fig. 5E). We applied this method to the LFP signal pretreated with a wide bandpass (5-40 Hz; 8th-order zero-phase Butterworth filter).

Fig. 5F shows snapshots of the spatial variation of the phase throughout the time course of a single SWD. Here, we observed complex traveling wave behavior that constantly shifted in direction and velocity, even across the duration of a single spike wave discharge. Fig. S6 shows a video of the phase evolution across multiple SWDs. Seizure propagation patterns were strikingly different with time through the seizure, demonstrating for the first time that seizure dynamics in absence epilepsy in this model do not have constant propagation trajectories or site of onset. More studies must be conducted to elucidate the origins of such variation. Importantly, our high channel density combined with a large area of recording along the cortical surface has allowed an unprecedented level of detail into seizure dynamics at micrometer-level precision.

## Discussion

We have introduced a novel system design and connectorization method, flex2chip, which increases the channel density of ultra-conformable, thin-film, flexible devices by 17 times in comparison to state-of-the-art devices. This enables multi-thousand channel counts on a single ASIC, unlike current approaches which rely on the modularization of multiple ASICs and circuit boards (*11, 19, 32, 33*). The assembly method is simple, consisting of placing the device on a CMOS-MEA with a thin layer of IPA. As the IPA dries, the microstructures self-assemble in their collapsed configuration. The connectorization does not require specialized equipment as needed for wire-bonding, anisotropic conductive film (*15*), or ultrasonic-on-bump bonding (*16*)).

Here, the key limitation to pad density is not the interconnection interfacial area but the routing of the traces. Although we demonstrate that we can form interconnections with a pitch of 50 µm with a theoretical density of 400 channels / mm^2^, the density for our 2200-channel devices is 272 channels / mm^2^ even with 1-µm-wide traces with 1 µm spacing. The input-referred noise level in the action potential range (300 Hz to 10 kHz) is 6.93 ± 3.30 µVrms, comparable to that of monolithic silicon probes such as the Neuropixels 2.0 (8.2 µV_rms_) (*34*).

The bonding methodology is agnostic to the chip, and rapid progress in their functionality, such as the addition of fast scan cyclic voltammetry, impedance spectroscopy, stimulation artifact suppression units, etc. can be facilely extended to the polymer device (*35*). Furthermore, it is also agnostic to downstream geometry, as shown here with both acute brain slice and in vivo cortical recordings. The developed technology can be easily used to extend the work on cutting-edge chronic intracortical and organoid recordings with mesh electronics (*36*–*38*). Improvements to miniaturize the CMOS-MEA onto a headstage for freely behaving mice experiments are already on the way.

## Materials and Methods

### flex2chip device fabrication

The devices were fabricated using standard microfabrication processes. The fabrication steps are as follows: (i) A 4-inch silicon wafer (P-type silicon, 0.1 – 0.9 Ohm cm; Silicon Valley Microelectronics Inc., CA, USA) was cleaned with O_2_ plasma (27.6 sccm, 50 W, 300 mTorr, 1 minutes; PE II-A, Technics, CA, USA). (ii) Alignment marks were patterned on the wafer. First, the wafer was dehydrated at 150 ºC and primed with hexamethyldisilazane (HMDS) (YES LP-III, Yield Engineering Systems, USA). Second, a 0.7-μm-thick layer of positive photoresist (I Line SPR 955 CM-0.7, Dow, MI, USA) was spin-coated on the wafer (1700 rpm, 30 seconds), and baked (90 ºC, 120 seconds). Third, the alignment marks were exposed (150 mW/cm^2^; PAS 5500/60, ASML, Veldhoven, Netherlands). Fourth, post exposure the resist was baked (90 seconds), developed with MF-26A (60 seconds, Dow, MI, USA), rinsed with water, and hard-baked (110 ºC, 60 seconds). Fifth, the alignment marks were etched 120 nm deep into SiO_2_ (100 sccm CF_4_, 2 sccm O_2_, 500 W, 250 mTorr, 40 seconds; P5000, Applied Materials, CA, USA). Finally, the photoresist was stripped using a microwave plasma system (LoLamp, 45 seconds; Aura Asher, Gasonics, CA, USA). (iii) A 250-nm-thick Ni sacrificial layer was deposited with electron beam evaporation (2.5 Å/s, 6e-7 Torr; ES26C, Innotec, MI, USA). (iv) A 1-μm-thick layer of polyimide (PI2610, Dupont, DE, USA) forming the substrate base was spin-coated (3000 rpm, 60 seconds), soft-baked (90 ºC, 3 min), and finally hard-baked in an inert N_2_ atmosphere (325 ºC, 30 min, 2 ºC/min ramp; Blue-M, PA, USA). (v) A metallic layer which forms the electrodes, interconnects, and bonding pads was then deposited. First, a 200-nm-thick liftoff layer (Microposit LOL 2000, Dow, MI, USA) was spin-coated (3000 rpm, 60 seconds) and baked (200 ºC, 7 minutes). Second, positive photoresist was patterned as described above. Third, the photoresist was descummed with O_2_ plasma (27.6 sccm, 50 W, 300 mTorr, 2.5 minutes; PE II-A, Technics, CA, USA). Fourth, a 10-nm-thick Cr and 100-nm-thick Pt layer were deposited with electron beam evaporation (1 Å/s, 6e-7 Torr; ES26C, Innotec, MI, USA). Finally, the photoresist was lifted off overnight (Microposit Remover 1165, Dow, MI, USA). (vi) A 1-μm-thick layer of polyimide which formed the insulating layer was spin-coated as described above. (vii) A negative mask, which defined the shape of the device as well the exposure of the electrode pads was deposited. First, positive photoresist was patterned and descummed as described above. Second, a 50-nm-thick Ni layer was deposited with electron beam evaporation (1 Å/s, 6e-7 Torr; ES26C, Innotec, MI, USA). Third, the photoresist was lifted off as described above. Fourth, the unprotected polyimide was then etched with O_2_ plasma (60 sccm O_2_, 400 W, 200 mTorr, 100 seconds; P5000, Applied Materials, CA, USA). (viii) Finally, the Si wafer was transferred to a Ni etchant solution (40% FeCl3:39% HCl:H2O=1:1:20) to remove the sacrificial Ni layer and negative mask and to release the device from the Si wafer.

### CMOS MEA modifications

The CMOS-MEA (Maxwell Biosystems, Zurich, Switzerland) has 26,400 pixels, 1024 of which can be arbitrarily chosen to record from simultaneously at 20 kHz (*23*). The bare die was wire-bonded to custom PCBs (3100 Plus, ESEC, Cham, Switzerland). The wirebonds were then encapsulated with epoxy (353ND-T, Epoxy Technology, MA, USA). Here, the pixels were recessed by a 1.2-μm-thick layer of SiO_2_ and SiN_x_. To elevate the pixel, Au was electrochemically deposited (60 seconds, 1.5 V, NB Semiplate AU 100 AS, Microchemicals, Ulm, Germany) using a Pt counter electrode.

### flex2chip device assembly

For the device assembly it is critical to minimize the number of particles to maximize the connectivity yield. The assembly of the device onto the CMOS MEA is as follows. (i) The CMOS MEA was transferred to a bath of sub-micron filtered IPA (Millipore Sigma, MA, USA, PX1838), cleaned by ultra-sonication for 30 seconds, and blown dry. (ii) A thin layer of IPA was applied using a pipette on the surface of the CMOS MEA as the lubrication layer, and the device was then placed on top with a pair of paintbrushes. (iii) The device was then sterilized in 70% ethanol solution for 1 hour and exposed to UV for 3 minutes.

Although Ohmic connectivity has already been established through the procedure above, the connection interface is only held with van der Waals forces and is delicate to external mechanical forces. To further secure it, a PDMS (Sylgard 184, Dow, MI, USA) block can optionally be placed on top of the flexible device and pressed down using a custom designed 3D-printed piece.

### Electrical characterization

Impedance and I-V curve measurements were conducted on a CMOS-MEA phantom using a potentiostat (SP-200, BioLogic, Seyssinet-Pariset, France). The distal end of the device was shorted to a gallium droplet to ensure Ohmic contact to probe the flex2chip interface at the proximal end. Connectivity, noise and signal attenuation measurements were conducted on the CMOS-MEA, as described by previous work (*23*). The distal end of the device was immersed in a PBS bath (ThermoFisher, MA, USA, 10010-023) along with a Pt counter electrode. For connectivity and signal attenuation measurements, the gain of the amplifiers was set to 24, and a 1-kHz 5-mV sine wave was injected at the counter electrode. Pixels connected to flex2chip microstructures would then record the sinusoidal waveform, and the corresponding pad could then be mapped. The attenuation was calculated to be the ratio between the measured and injected signal. Pt-black coating (100 mM hexachloroplatinic acid, Millipore Sigma, MA, USA, 206083) was electrochemically deposited (0.5 V, 30 seconds) under mechanical agitation to lower the electrode-electrolyte impedance

### Brain Slice Preparation

All use of mice and experimental protocols was approved by the Basel Stadt veterinary office according to Swiss federal laws on animal welfare. Wild type mice (postnatal day 14, C57BL/6JRj, Janvier Labs) were decapitated under isoflurane anesthesia and the brains were removed and immersed into ice-cold carbogen-bubbled (95% O2 + 5% CO2) ACSF solution containing (in mM): 125 NaCl, 2.5 KCl, 25 glucose, 1.25 NaH_2_PO_4_, 25 NaHCO_3_, 2 CaCl_2_, 1 MgCl_2_. The cerebellum was dissected and glued on the cutting stage of a vibratome (VT1200S, Leica, Wetzlar, Germany). Sagittal cerebellar slices of 380 µm thickness were obtained. Slices were then maintained in ACSF at room temperature (RT) until use.

A 6 mm diameter stainless-steel ring was bonded to a 720-channel device using cyanoacrylate adhesive (Pattex, Henkel, Aachen, Germany), where the exposed platinum electrodes were facing away from the ring. The construct was placed on the acute slice and held in place with the stainless steel ring. The tissue was continuously perfused with carbogen-bubbled ACSF at 33-36 °C to maintain cell viability.

### In vivo preparation

Mice with the heterozygous loss of function mutation in *Scn8a* (male, 12 weeks old, C3HeB/FeJ-Scn8amed/J, Jackson Laboratory, Stock#: 003798 (*39*)), referred to in this manuscript as *Scn8a*^+/-^, were the vertebrate animal subjects used for in vivo measurements. All procedures performed on the mice were approved by Stanford University’s Administrative Panel on Laboratory Animal Care (APLAC, Protocol #12363). The animal care and use programs at Stanford University meet the requirements of all federal and state regulations governing the humane care and use of laboratory animals, including the United States Department of Agriculture (USDA) Animal Welfare Act, and the Public Health Service (PHS) Policy on Humane Care and Use of Laboratory Animals. The laboratory animal care program at Stanford is accredited by the Association for the Assessment and Accreditation of Laboratory Animal Care (AAALAC). All mice were maintained on a reverse 12-hour dark/light cycle (temperature: 20-25 ºC, humidity: 50-65 %) in the Stanford University’s Veterinary Service Center (VSC) and fed with food and water ad libitum as appropriate. All experiments occurred during their active cycle.

### Surgery

Anesthesia was induced with isoflurane (4%; maintained at 1.5%) followed by injection of carprofen (2 mg/kg). Fiducial marks were marked on the skull at the following coordinates from bregma: anteroposterior, -0.85 mm; mediolateral, 2.5 mm. A self-tapping bone screw (Fine Science Tools, CA, USA, 19010-10) was set in the skull over the cerebellum. A stainless steel headbar was cemented onto the skull using dental cement (C&B Metabond, Parkell, NY, USA). After headbar implantation, mice were habituated to run on a treadmill for 7 days. In a second surgical procedure, a 2-mm diameter craniotomy was then made over the fiducial mark and covered with KwikCast (World Precision Instruments, FL, USA). The mouse then recovered overnight before the recording session.

### Electrophysiological recording

Once mounted on the treadmill, the KwikCast above the craniotomy was removed and the well was filled with saline (0.9% NaCl). The ECoG device was laid on the surface of the cortex and the saline was wicked away to allow sufficient contact between the device and cortex. KwikCast was then reapplied. The CMOS-MEA counter electrode operates at a reference voltage of 1.65 V. Consequently, the animal was isolated from ground by connecting the reference of the chip to a skull screw. Recording then lasted for 40 minutes.

### Spike sorting

Data analysis was performed using custom software written in Python 3.9.0 and MATLAB 2019b (Mathworks, MA, USA). Automatic spike sorting was performed using Kilosort 3 (https://github.com/MouseLand/Kilosort) (*34, 40*). Subsequently, using the ecephys spike sorting pipeline (https://github.com/AllenInstitute/ecephys_spike_sorting), double counted spikes were removed from each cluster (within ±0.16 ms), and the ISI violations within the refractory period (± 1.5 ms) were calculated. Units were only classified as good if the number of spikes was greater than 100, the ISI violations were less than 0.1, and if Kilosort 3 originally labelled the spike as ‘good’. Finally, clusters were inspected and curated in Phy (https://github.com/cortex-lab/phy).

## Author Contributions

E.T.Z and N.A.M. conceived the experiments. E.T.Z. and P.W. fabricated the devices. E.T.Z. and A.P. developed the assembly process. E.T.Z., N.H., H.U. and A.Z. characterized the devices. H.U., S.R. and E.T.Z. prepared the CMOS-MEAs. J.H. and E.T.Z. performed experiments in rodents and analyzed the data. J.B. and E.T.Z. performed experiments on cerebellar slices and analyzed the data. N.A.M., A.H., and J.R.H. supervised the work. E.T.Z. and N.A.M. wrote the manuscript with contribution from all the authors.

## Data and materials availability

All data needed to evaluate the conclusions in the paper are present in the paper and/or the Supplementary Materials. Additional data related to this paper may be requested from the authors.

## Supplementary Materials

**Figure S1.**
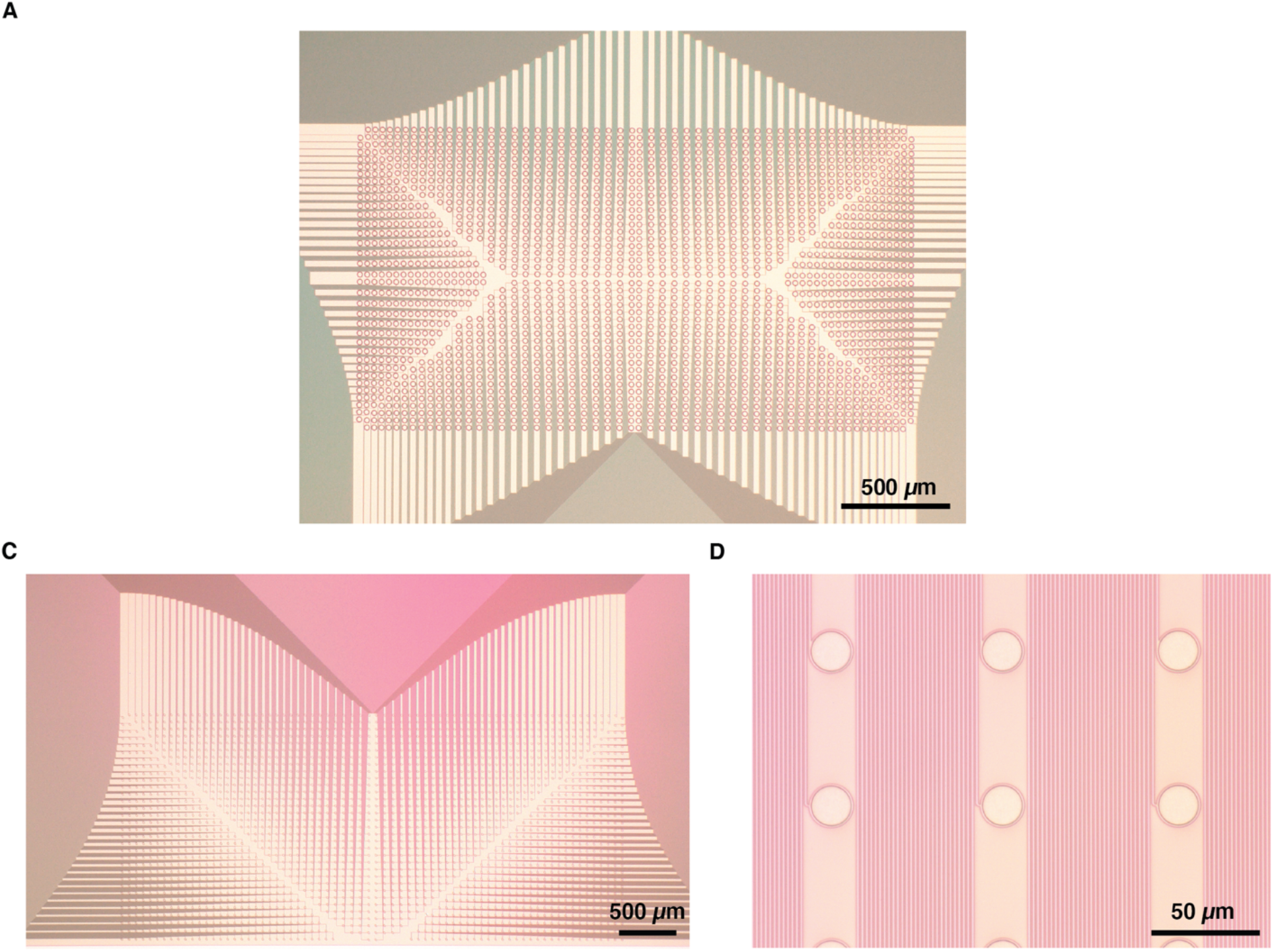
2200-channel geometry. (**A**) Microscopy image of 2200 flex2chip microstructures in a 3.85 mm x 2.10 mm area, with a channel density of 272 channels / mm^2^. The routing escapes in all four directions to maximize the density of pads in the active area. (**B**) Microscopy image of the distal end of the 2200-channel device, which interfaces with the neurons. (**C**) An inset of Fig. S1B showing each individual recording/stimulation pad.

**Figure S2.**
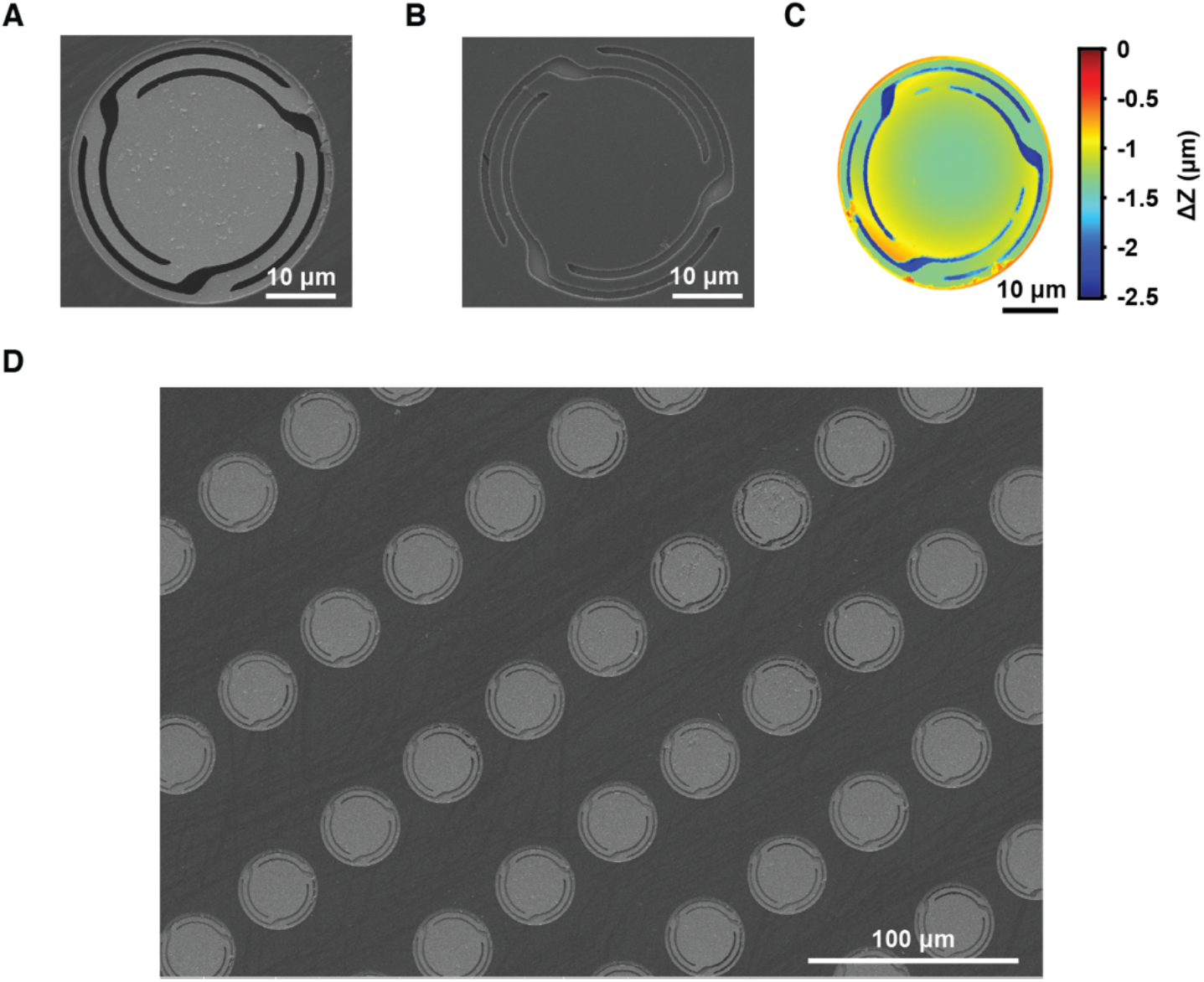
flex2chip microstructure characterization. (**A** and **B**) SEM images of the flex2chip microstructure, platinum-side facing up and down respectively. Here, the devices were released from the wafer and placed on a glass slide. (**C**) Optical profilometer image of the flex2chip microstructure, platinum-side facing up. (**D**) SEM image of an array of flex2chip microstructures, platinum-side facing up.

**Figure S3.**
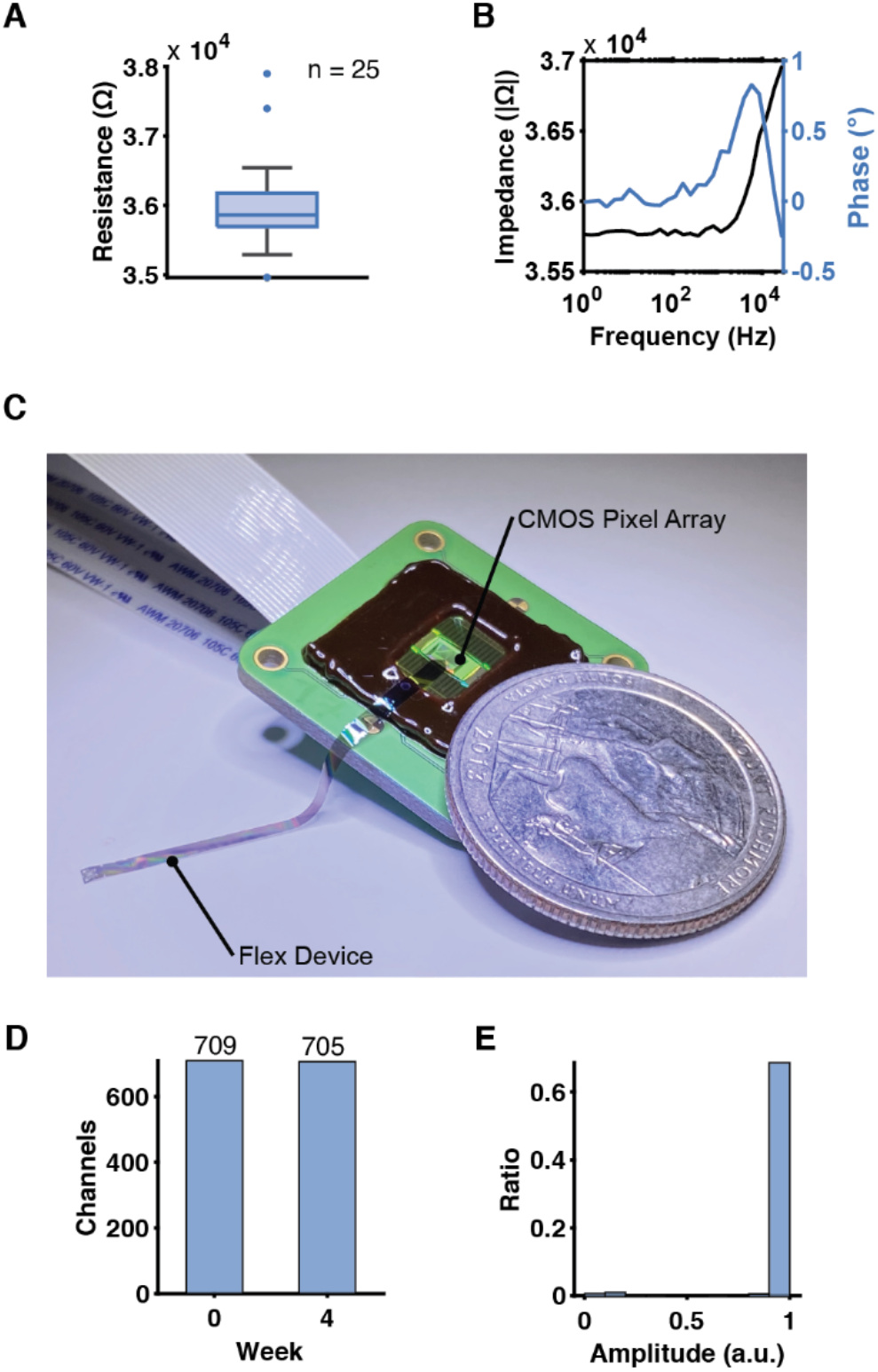
flex2chip electrical characterization. (**A**) The resistance of the flex2chip interface computed from the IV curves from Fig. 2D. The narrow variation (36.0 ± 0.60 kΩ, n = 25) in impedance demonstrates minimal interelectrode variability. (**B**) A representative Bode plot indicates uniform resistivity and phase from 1 Hz to 30 kHz. This finding confirms that there is no capacitive impedance at the physiological frequencies of interest. (**C**) Photograph of an assembled flex2chip device on a CMOS-MEA adjacent to a United States quarter. The CMOS-MEA headstage has two 24-channel ZIF connectors leading to a downstream FPGA. (**D**) The connectivity yield is virtually unchanged after leaving it in an incubator at 37 °C and 97% humidity over a period of a month, with a minor decrease of 709 to 705 out of a total of 720 pads. (**E**) The electrical characteristics also remain unchanged, with the aged device detecting 0.99 ± 0.03 of the full-scale signal.

**Figure S4.**
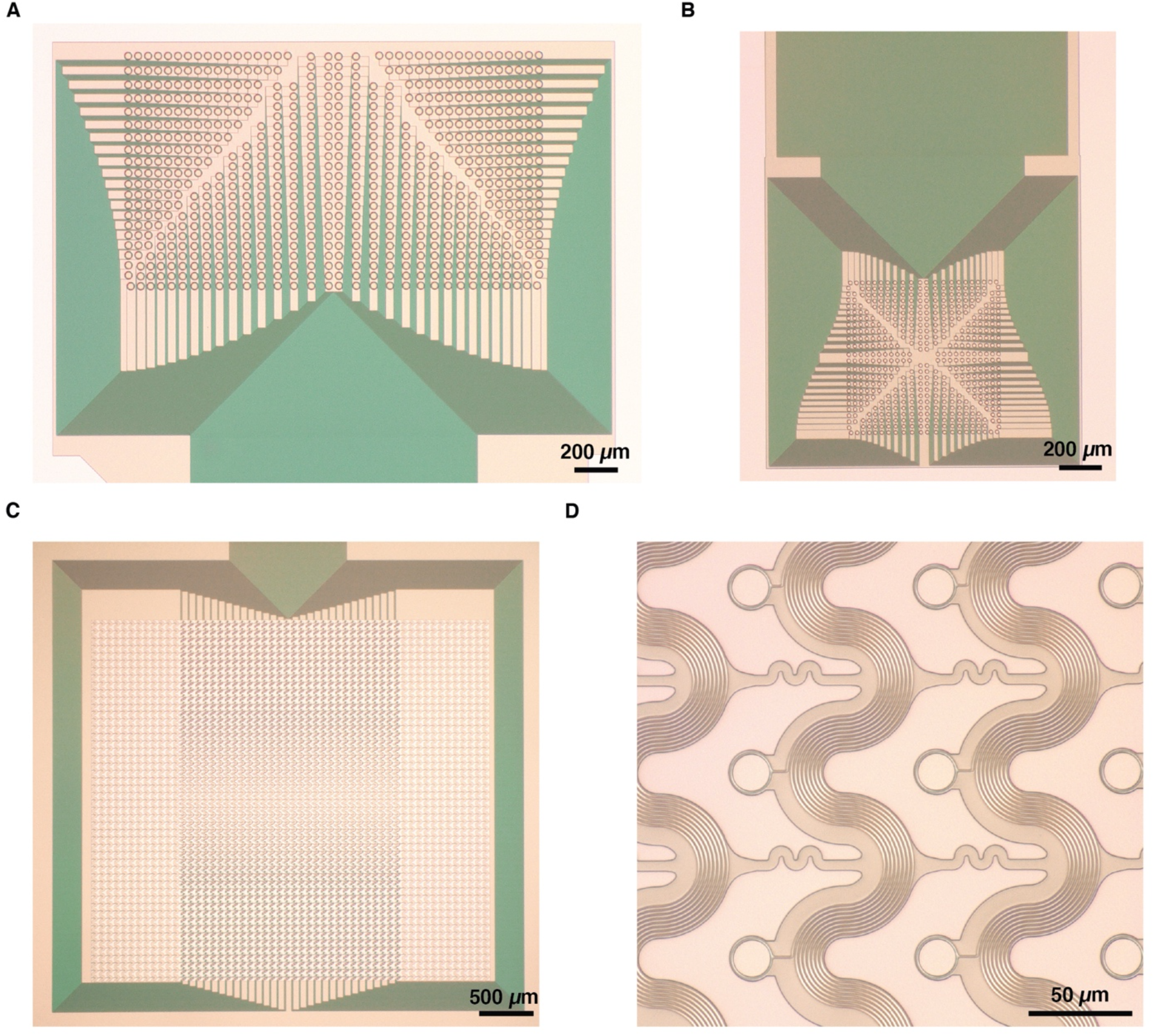
720-channel geometry. (**A**) Microscopy image of 720 flex2chip microstructures in a 2.10 mm x 1.20 mm area, with a channel density of 286 channels / mm^2^. This proximal end of the device interfaces with the CMOS-MEA to establish low-noise and Ohmic contact. Each microstructure has an individual 1-µm-wide trace which is routed to the distal end of the device. (**B** and **C**) The distal end of the device can be freely tailored depending on the physiological system in question. The first is an ECoG grid, with 504 20-µm-diameter recording pads in a 0.76 mm x 0.76 mm area, with 216 open channels. The device was designed to fit within a 2-mm-diameter craniotomy for awake head-fixed recordings. The second is a fenestrated grid of 720 20-µm-diameter recording pads in a 2.63 mm x 2.09 mm area. This design was used for brain slice experiments where the fenestrations allow nutrient and oxygen diffusion in the slice during the recording sessions. (**D**) An inset of Fig. S4C showing the individual recording pads.

**Figure S5.**
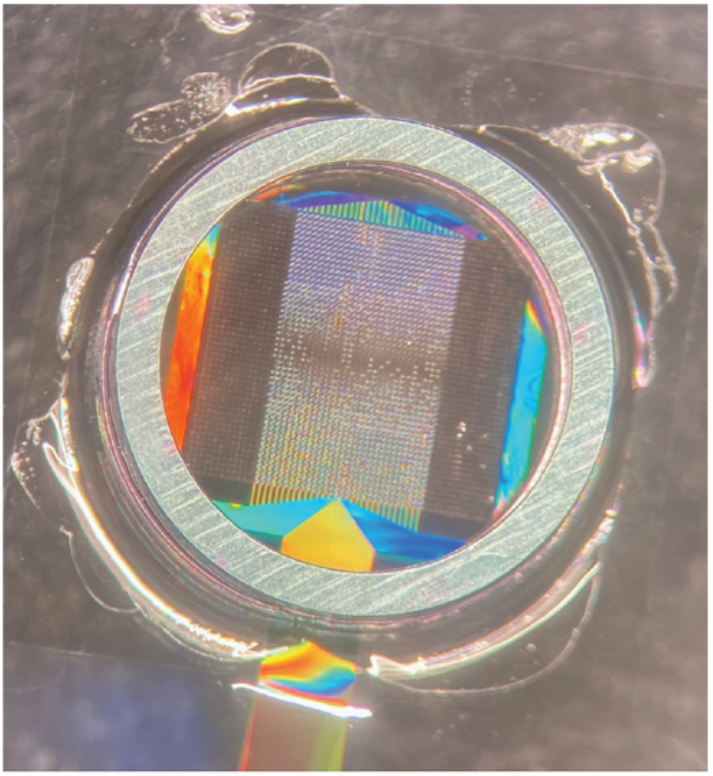
In vitro device setup. The 720-channel device has an active area of 2.63 mm x 2.09 mm with electrodes at a 90 µm pitch. A 6-mm-diameter stainless-steel ring was attached to the device using superglue so that sufficient mechanical contact was made when placed on top of the slice. The fenestrations allow for nutrient and oxygen diffusion so that the slice remains healthy for the duration of the experiment.

